# Structure and function of diadenylate cyclase DacM from *Mycoplasma ovipneumoniae*

**DOI:** 10.1101/2022.03.11.483894

**Authors:** Xiujing Hao, Xinhui Zhou, Ying Zhang, Yang Han, Zhaokun Xu, Chunji Ma, Haixia Luo, Kemin Qi, Shilong Fan, Min Li

## Abstract

Cyclic diadenosine monophosphate (c-di-AMP) is a second-messenger nucleotide that is produced by many bacteria. C-di-AMP can not only regulate bacterial growth, cell-wall homeostasis, ion transport and gene transcription, but can also be recognized by multiple sensor / receptor proteins in infected host cells to trigger an innate immune response. *Mycoplasma ovipneumoniae* causes non-progressive pneumonia in both sheep and goats. Here, we analyzed c-di-AMP signaling in *M. ovipneumoniae*, which is a genome-reduced obligately pathogenic bacterium. Our results demonstrate that these bacteria can produce c-di-AMP, and we could identify the diadenylate cyclase, which was named DacM. The enzyme was found to utilize both ATP and ADP to synthesize c-di-AMP, resembling CdaM from a novel family of diadenylate cyclases first found in *Mycoplasma pneumoniae*. Furthermore, we present the crystal structures of DacM in the apo state and substrate-bound state at 3 Å and 1.9 Å resolution, respectively. Mutation of residues Asp112, Gly113, Tyr128, Phe129, and Arg143 surrounding the active sites to Ala were lethal to DacM enzymatic activity. These structures provide valuable insights into the biochemistry of c-di-AMP, and offer a basis for the structure-based design of new drugs for animal husbandry.

## Introduction

Mycoplasmas are distinguished from other bacteria by minute size and lack of a cell wall[1]. They have been studied as some of the smallest free-living and self-replicating cells[2]. *Mycoplasma ovipneumoniae* infects sheep and goats, causing pleuropneumonia. The clinical signs of pleuropneumonia include lethargy, chronic coughing, rectal prolapse, and poor weight gain[3]. However, there is a lack of specific drugs and vaccines for the treatment and prevention of *M. ovipneumoniae*.

Cyclic diadenylate monophosphate (c-di-AMP) is an important second messenger, and its function in bacteria has been gradually clarified in recent years. It plays important roles in microbial growth[4, 5], DNA integrity[6], balance of cell wall metabolism[7, 8], biofilm formation[9], transport of potassium ions[10], resistance to abiotic stress[11], synthesis of fatty acids[12], virulence[4, 9] and antibiotic resistance[5]. These studies show that c-di-AMP participates a variety of cellular processes in bacteria, with distinctive roles in different species[13]. In addition, c-di-AMP also plays a key role in activating innate immunity during host-microbe interactions[14-16]. Additionally, excessive intracellular amounts of c-di-AMP can become toxic[17].

C-di-AMP is synthesized by diadenylate cyclases (DACs) using two molecules of ATP, or in some cases two molecules of ADP[18, 19]. All known DACs possess the conserved diadenylate cyclase domain (DisA_N domain), which catalyzes the cyclase reaction in c-di-AMP synthesis. The DisA_N domain contains conserved DGA (Asp-Gly-Ala) and RHR (Arg-His-Arg) motifs[20, 21]. DisA_N domain proteins have been found in many bacteria and archaea. The importance of c-di-AMP for the growth of several pathogenic bacteria is marked by an increased resistance to antibiotics that target the cell wall. Moreover, the lack of DAC enzymes in humans makes them an ideal target for the development of novel antibiotics. Currently, five types of DACs have been identified, named DisA, CdaA (or DacA), CdaS, CdaM and CdaZ[19, 22]. Most bacteria contain a single DAC enzyme, some contain multiple enzymes, such as Clostridium spp., which contains two types of DAC (CdaA and DisA) and B. subtilis, which contains DisA, CdaA, and CdaS[13, 14]. However, the detailed molecular mechanism of c-di-AMP regulation is still unknown. Thus, more structural and functional investigations of DAC enzymes are required for a better understanding of c-di-AMP synthesis urgently.

The structures of three types of DAC enzymes have been determined to date, including the crystal structure of DisA from Thermotoga maritima[23], CdaA from Listeria monocytogenes[17, 24], DacA from Staphylococcus aureus[7] and CdaS from Bacillus cereus[19]. Among them, DisA from Thermotoga maritima was the first protein with a DisA_N domain to be characterized[23]. The structure showed the enzyme in a homo-octameric state with two adjacently positioned DAC domains, each binding one ATP molecule, catalyzing the synthesis of c-di-AMP[6]. The structure of CdaA from Listeria monocytogenes revealed the flexibility of a tyrosine side chain involved in locking the adenine ring after ATP binding. The essential role of this tyrosine was confirmed by the drastic loss of enzymatic activity following its mutation to Ala[17]. However, there is still no structural information for CdaM and CdaZ-type DACs.

In this study, we discovered c-di-AMP synthesis in *M. ovipneumoniae* strain Y98, and identified the diadenylate cyclase, which we named DacM. This novel c-di-AMP synthase was found to only contain a single transmembrane domain, indicating that it belongs to the CdaM-type. Although the protein also contains conserved RHR (Arg-His-Arg) and DGA (Asp-Gly-Ala) motifs, the overall sequence has low similarity with the reported c-di-AMP synthase sequences. Furthermore, we solved the crystal structures of DacM in the apo state and substrate-bound state, which could facilitate the design of diadenylate cyclase inhibitors, with great potential for applications as novel antibiotics in animal husbandry.

## Results

### *M. ovipneumoniae* can produce c-di-AMP and its genome encodes potential c-di-AMP cyclases

To determine whether *M. ovipneumoniae* Queensland Strain Y98 can produce c-di-AMP, nucleotides were extracted and analyzed using an ACQUITY UPLC I-Class system coupled with a VION IMS QTOF mass spectrometer. The retention time (tr) of synthetic c-di-AMP was 5.71 min under these conditions (Fig 1B). A peak with the same retention time was observed in the extract of *M. ovipneumoniae* Y98 cells (Fig 1A).

**Fig 1.**
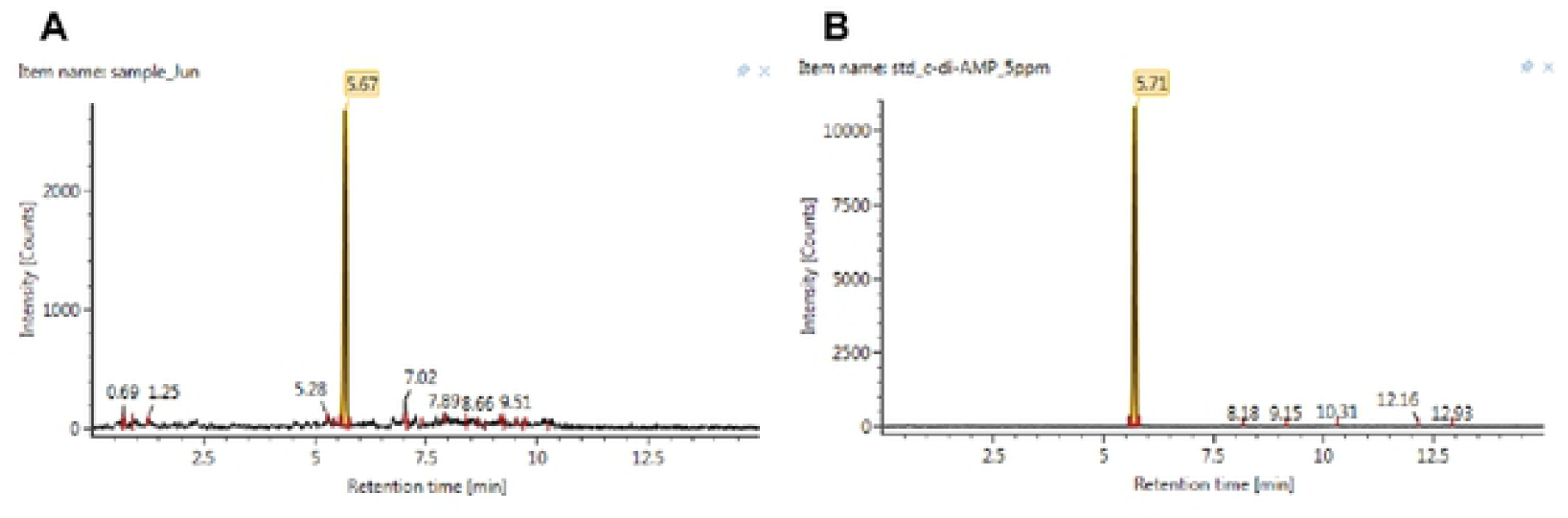
Detection of c-di-AMP in nucleotide extracts from *M. ovipneumoniae Y98* by LC-MS analysis. The peak at 5.67 min represents c-di-AMP from *M. ovipneumoniae Y98* (A), as determined from its correspondence to the retention time of chemosynthetic c-di-AMP (B).

To identify potential enzymes involved in c-di-AMP synthesis in *M. ovipneumoniae* Y98, we searched for potential proteins with a DAC domain using a whole-genome sequencing approach. We found a single protein containing a DAC domain, with conserved ‘DGA’ (Asp-Gly-Ala) and ‘RHR’ (Arg-His-Arg) residues. We named it DacM (diadenylate cyclase of *M. ovipneumoniae*). It shares the highest similarity to the diadenylate cyclase from *M. pneumoniae* (CdaM) and the N-terminal domain of this protein has only one transmembrane helix, as well. The domain arrangement of DacM is shown in S1 Fig.

### DacM is a diadenylate cyclase that can use both ATP and ADP as substrates

In order to test whether DacM does indeed exhibit diadenylate cyclase activity, we took advantage of the inability of *E. coli* to produce c-di-AMP. The c-di-AMP produced by *E. coli* BL21 (DE3) carrying the corresponding pET28b-*dacm* plasmid was detected. The same strain carrying the empty vector pET28b was used as a negative control. As reported previously[25], no c-di-AMP was detected in the strain with the empty vector, while the *E. coli* BL21 (DE3) strain with the pET28b-*dacm* vector produced detectable levels of c-di-AMP. This result demonstrated the diadenylate cyclase activity of DacM (Fig 2A).

**Fig 2.**
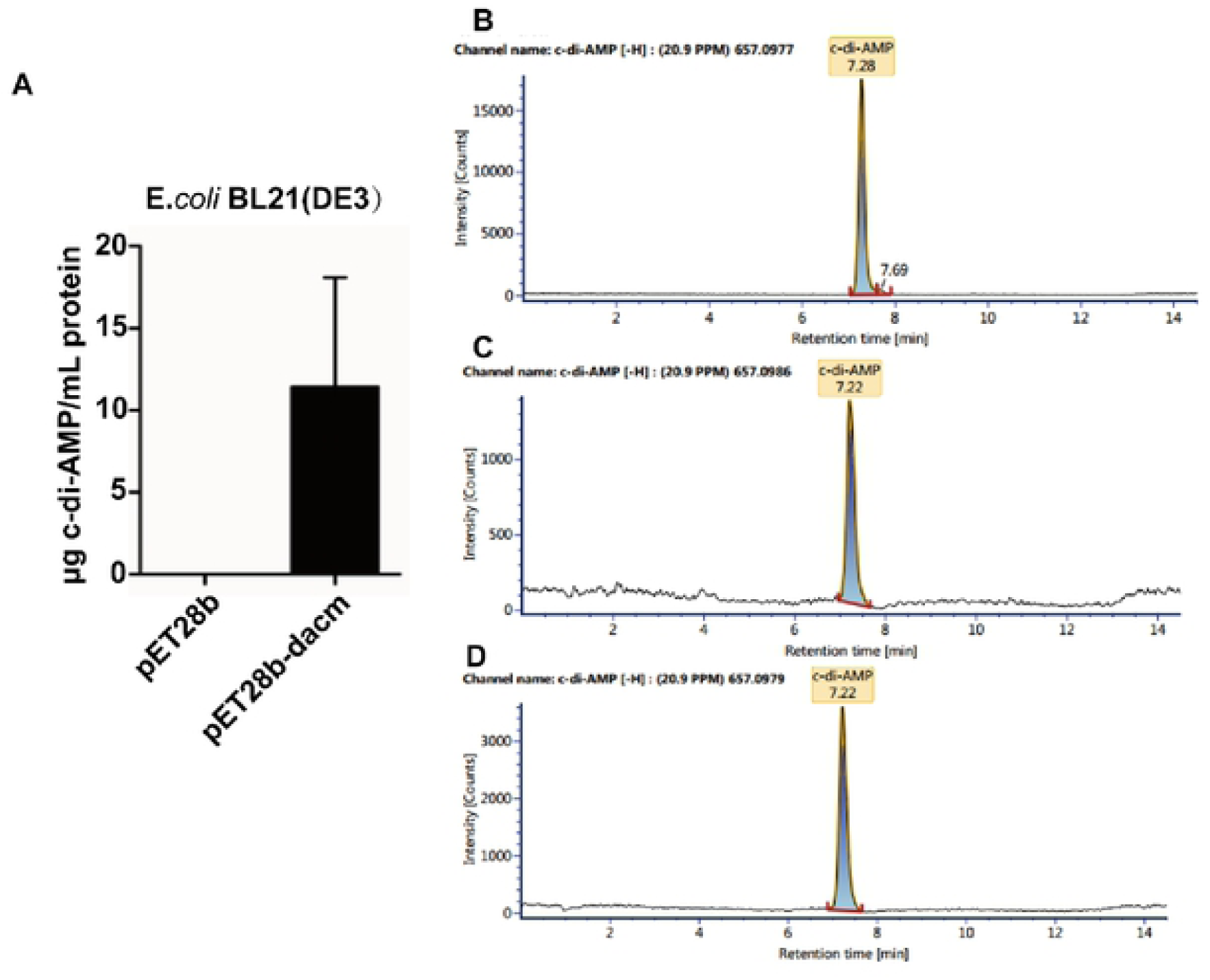
Enzymatic activity of DacM in vivo and vitro. (A) C-di-AMP concentration in cells of E. coli BL21(DE3) carrying the empty vector pET28b or the DacM expression vector pET28b-dacm. The c-di-AMP concentrations were determined by LC-MS analysis, and the results show the average and standard deviation of three values plotted as μg c-di-AMP/ml. (B-D) Enzymatic activity of DacM in vitro with ATP and ADP as substrates. (B) C-di-AMP standard. (C) ATP as substrate in the enzymatic reaction system. (D) ADP as substrate in the enzymatic reaction system.

To study the enzymatic properties of DacM in vitro, recombinant N-terminal His_6_-fusion protein was purified from *E. coli* (S2 Fig). The DAC activity and substrate spectrum of the protein were determined by conducting in vitro enzyme activity assays and characterizing the products by LC-MS. The c-di-AMP in the reaction mixture was identified by comparing the retention time of the product with commercially available c-di-AMP. The results indicated that DacM possesses diadenylate cyclase activity and can convert both ATP and ADP into c-di-AMP (Fig 2B-D).

### Localization of diadenylate cyclase DacM

DacM was predicted to contain a transmembrane helix, but there were no reports of its intracellular localization in vivo. In order to investigate the localization of DacM, immunogold staining and transmission electron microscope (TEM) imaging were performed. Based on the observations of DacM in TEM micrographs, most *M. ovipneumoniae* presented circular morphology with immunogold staining mainly localized around the cell perimeter after incubation with the anti-DacM antibody. These observations indicated that DacM is a membrane-associated protein (S3 Fig).

To further verify whether DacM is a membrane-associated protein, membrane proteins of *M. ovipneumoniae* Y98 were extracted using 1.5% (v/v) Triton X-114, and subjected to western blot analysis. The results confirmed the presence of DacM in the extracted membrane protein fraction (S4 Fig).

### Overall Structure and Oligomerization of Apo DacM

A truncated version of DacM (Δ^2-33^DacM), which lacks the transmembrane helix, was used for the crystallography trials. Fortunately, the crystal structure of the apo state of DacM from *M. ovipneumoniae* Y98 could be determined at a resolution of 3 Å. The crystal belonged to space group I2_1_3 (S1 Table) and the phase problem was solved by molecular replacement using the structure of CdaA from *Listeria monocytogenes* (PDB ID: 4RV7) as the search model. Each asymmetric unit in the crystal contains two DacM protomers. The folding and orientation of the two protomers was very similar, and they superimposed very well with an rmsd of 0.362 Å, but a few different features were seen. The major difference was observed in the loop between β3 and α4 helix, and also near the N-terminal end of α4 helix (S5A Fig). Clearly, this loop region in protomer A flanks one side, while in protomer B this loop blocks access to the substrate cavity. In general, the overall structure of DacM showed a fold similar to its homologues, such as DacM or CdaA. However, it was composed of 5α helices surrounded by 6 mixed-parallel and anti-parallel β-strands in each protomer (Figs 3A and 3B). The interface between two protomers was surrounded by residues of helix 3 and the loop connecting helix 2 and helix 3 (Fig 3C). Further analysis revealed that two patches are involved in the stabilization of the homodimer. The contact in one patch is mediated by hydrogen bonds formed by residues Ile91 and Asp92 from one protomer with residues Leu94 and Arg121 from the second protomer, respectively. The contacts in the other patch are mediated by hydrogen bonds formed by residues Ala102 and Gln105 from one protomer with residues Ser108 and Ser107 from the second protomer, respectively.

**Fig 3.**
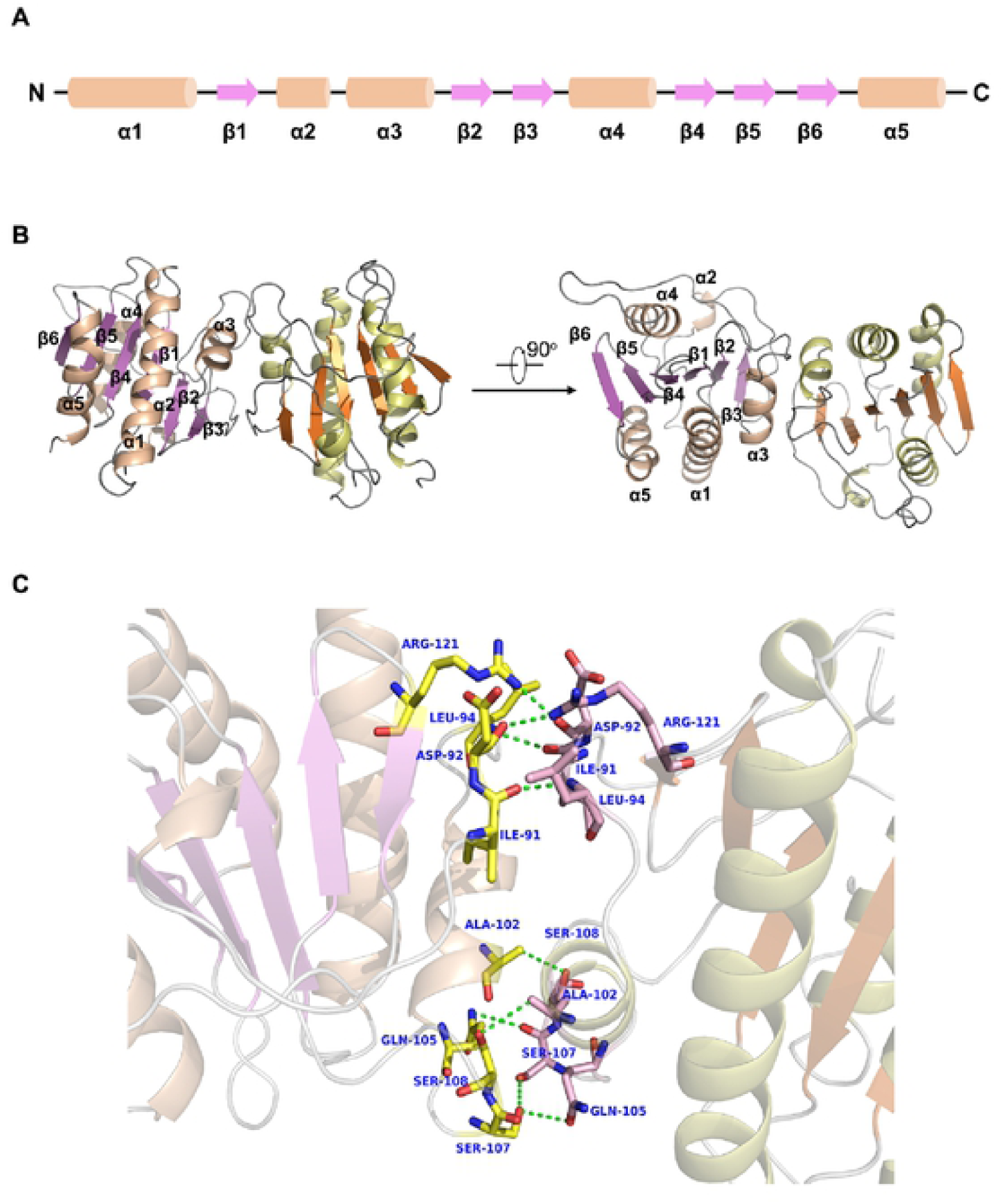
Crystal structure of DacM (Δ^2-33^DacM) in the apo state. (A) The secondary domain architecture of DacM (Δ^2-33^DacM) composed of five α-helixes and six β-strands. (B) Overall structure of the apo DacM (Δ^2-33^DacM) homodimer presented in two perpendicular views. Protomer A is shown with α-helixes in pink and β-strands in purple, and protomer B is shown with α-helixes in light green and β-strands in orange. (C) Close view of the dimer interface of DacM (Δ^2-33^DacM). The residues involved in the interaction within the homodimer are shown as sticks. Hydrogen bonds are indicated with green dashed lines.

### Structure of DacM bound to ATP / ADP and conformational changes between the apo and substrate-bound forms

To further investigate the enzymatic mechanism of DacM, we crystallized DacM (35-203) in the presence of ATP, ADP or c-di-AMP, respectively. To our surprise, we obtained crystals that diffracted very well in the presence of ATP as an additive. Finally, the structure of the substrate-bound form was refined to 1.9 Å. Overall, the solved structure also contained two protomers in each asymmetric unit, belonging to the P2_1_2_1_2_1_ space group. The two protomers in the asymmetric unit overlapped very well, with an rmsd of 0.228 Å (S5B Fig). Additionally, two other small electron density objects were found inside the structure. Further analysis reveals that one is consistent with an ATP molecule, which was clearly bound in the active site of one DacM protomer (protomer A) (Fig 4A). The other electron density, which was found in the active site of the second DacM protomer (protomer B), was consistent with a bound molecule of ADP, showing well-defined density of two phosphate groups (α- and β-phosphates) (Fig 4B). At the high resolution of the structure we obtained, it appears likely that protomer B catalyzed the ATP hydrolysis during the crystallization phase. A similar state was observed in a crystal of CdaA from *Listeria monocytogenes* previously described by Heidemann et al[17, 24], which simultaneously bound c-di-AMP and AMP in the homodimer.

**Fig 4.**
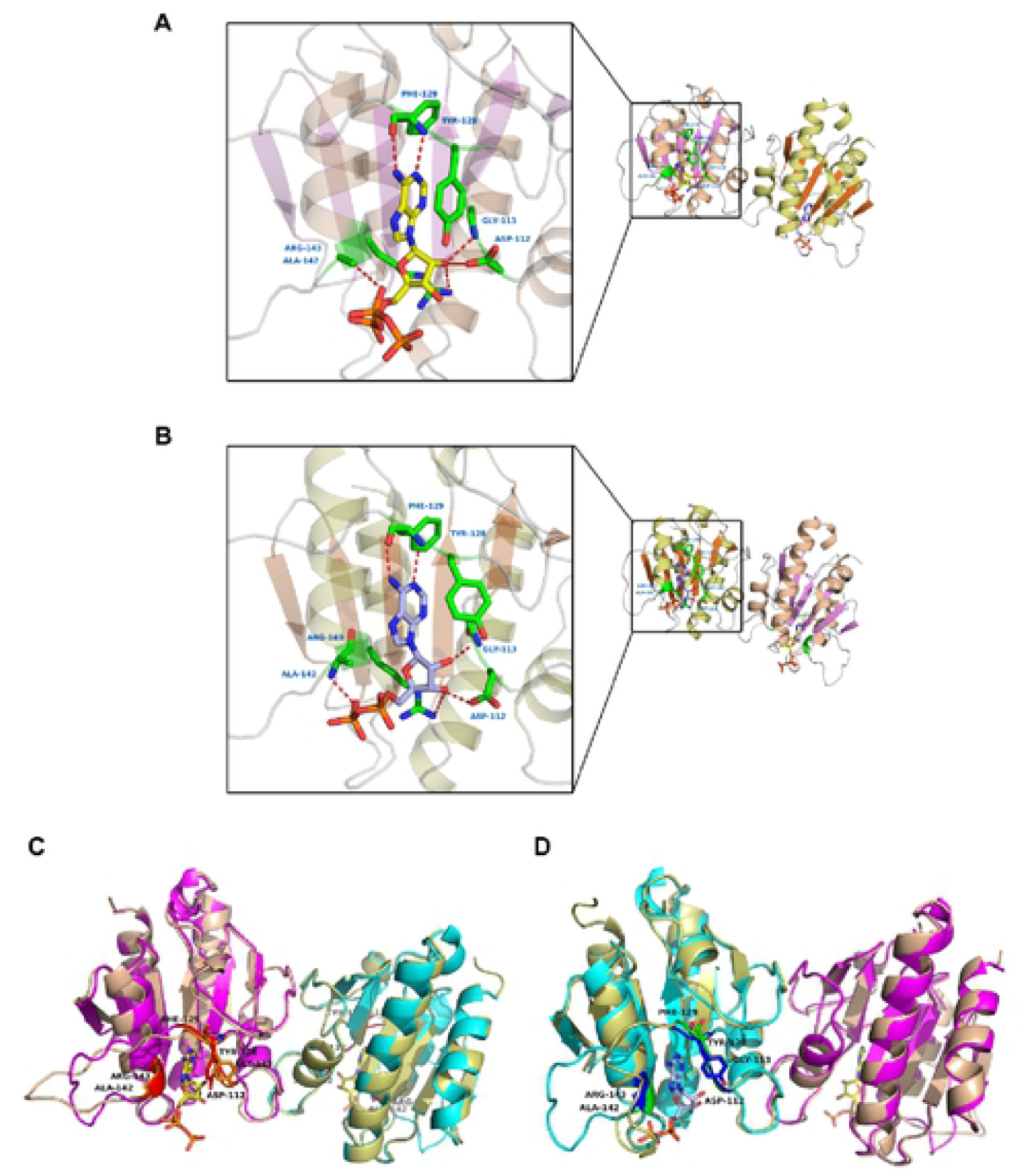
The active site of dimeric DacM with two different substrate molecules bound, and structural comparison between the apo form and substrate-bound form. (A) and (B), the overall shape of the substrate-bound form of DacM (Δ^2-33^DacM) and close views of the substrate binding pocket, i.e. ATP binding pocket (A) and ADP binding pocket. (B). Key residues involved in the active site are labeled and shown as sticks. The interactions between the substrate and residues are shown in red dashed lines. The main backbone of the ATP molecule is shown in yellow and the main backbone of the ADP molecule is shown in blue. (C) and (D), the overall structure and substrate binding site comparison between the apo form and substrate-bound form. The apo form of DacM is colored pink and light green (representing protomer A and B, respectively), while the substrate-bound form of DacM is colored magenta and cyan (representing protomer A and B, respectively). In panel C, amino acids involved in the ATP-binding site of protomer A are labeled and colored orange (apo form) and red (substrate-bound form). In panel D, amino acids involved in the ADP-binding site of protomer B are labeled and colored green (apo form) and dark blue (substrate-bound form). Among them, the characteristic Tyr 128 is presented as sticks. The conformational switch of Tyr128 is seen during the substrate binding process, especially in protomer A.

The active sites between the two protomers of DacM were also a slightly different (Figs 4A and 4B). In the ATP-binding cavity, residue Tyr128 located in the loop connecting β3 and α4, which is highly conserved among different homologues, engaged in a π-π-stacking interaction with the adenine moiety of ATP, as was reported for CdaA[17]. Compared to the apo state, the side chain of residue Tyr128 underwent an obvious rotation toward the adenine moiety of ATP (Fig 4C). Furthermore, the neighboring residue Phe129 also interacted with the adenine moiety via its amino group and carbonyl group. Additionally, the highly conserved residues Asp112 and Gly113, which belong to the ^112^D^113^GA motif, as well as Arg143, which belongs to the ^143^RHR motif, were all involved in the interaction with the ribose moiety. Additionally, the Ala142 residue coordinated the α-phosphate of ATP (Fig 4A).

However, in the ADP-binding cavity of protomer B, the side chain of the highly conserved Tyr128 presented a similar orientation with the apo state (Fig 4D). Therefore, the cavity was wider than the ATP-binding cavity and more space was free for the ADP molecule to occupy (Fig 4B). The highly conserved residues Asp112 from the ^112^D^113^GA motif and Arg143 from the ^143^RHR motif both interacted with the 3’-OH of the ribose moiety, instead of the interaction with the 2’-OH observed in the ATP-binding state. Due to the lack of the strong π-π-stacking interaction of Tyr128 and changes in the binding of the ribose moiety to Asp112, the phosphate backbone of ADP was more flexible than that of ATP. Therefore, it allowed the phosphate backbone of ADP to move closer to the α4 helix due to the interaction between residue Ala142 and the β-phosphate of ADP (Fig 4B).

Overall, the structures of the apo form and the substrate-bound form displayed similar features and largely overlapped, with an rmsd of 0.519 Å. Few major rearrangements of the overall structures were observed, but two small notable conformational changes were indeed observed in the substrate binding cavity. One was the orientation of Tyr128 in protomer A, which was rotated close to the ATP molecule when binding the substrate (Fig 4C). This rotation trapped ATP in the cavity with stronger interactions. The other was the loop near the N-terminal end of the α4 helix in protomer B, which moved away when binding the substrate (Fig 4D). This offered open access for ADP approaching the binding cavity. Taken together, these two changes both facilitate substrate binding in the homodimer.

### DacM mutants decreased or completely abolished the catalytic activity

To test whether changes in the active sites of DacM affect the ability of DacM to synthesize c-di-AMP, we designed the following single-point mutations: D112A, G113A, Y128A, F129A, R143A, and five-point simultaneous mutations (MUT). Whether these mutants were able to synthesize c-di-AMP was analyzed by HPLC technique. Compared with the wild type (WT) strain, trace amounts of c-di-AMP were detected in D112A, G113A, Y128A, F129A mutants, and c-di-AMP was not detected in R143A and MUT mutants. It can be seen that most mutants exhibited decreased DacM activity or completely abolished the catalytic activity, indicating their importance in maintaining the DacM enzymatic activity (Fig 5).

**Fig 5.**
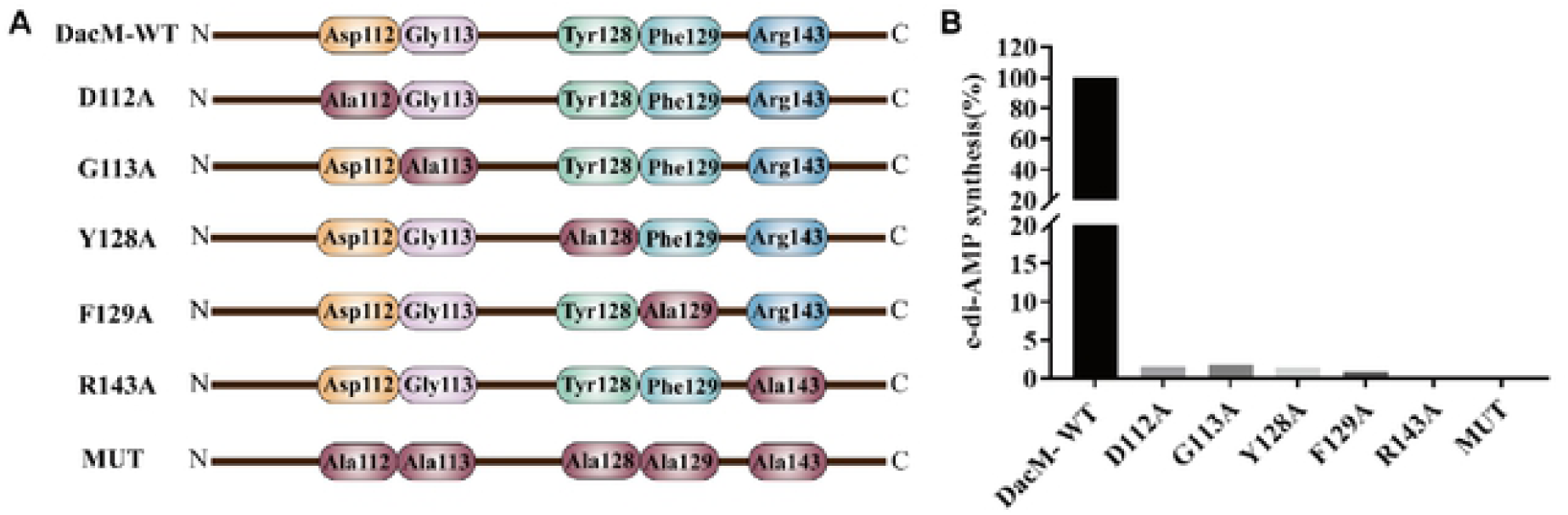
DacM mutants catalytic activity on c-di-AMP synthesis. (A) Schematic diagram of DacM mutants. (B) DacM mutants catalytic activity on c-di-AMP synthesis.

### Structural comparison between DacM and its homologues from other species

A structure-based sequence alignment between DacM and its homologues from other species, including CdaM from *Mycoplasma pneumoniae*, DacA from *Staphylococcus aureus*, CdaA from *Listeria monocytogenes*, CdaS from *Bacillus cereus* and DisA from *Thermotoga maritima*, is shown in Fig 6A. As expected, the residues we identified as being part of the active cavity in the structure were mostly highly conserved, with strict conservation of Asp112, Gly113 and Arg143 from the conserved motifs ^112^D^113^GA and ^143^RHR. Besides, the Tyr128 was conserved among DacM, CdaM, DacA and CdaA. This residue was reported to have an essential role for the enzymatic activity, but the variation of Tyr between CdaS and DisA suggests a potentially different mechanism of substrate binding in different species.

**Fig 6.**
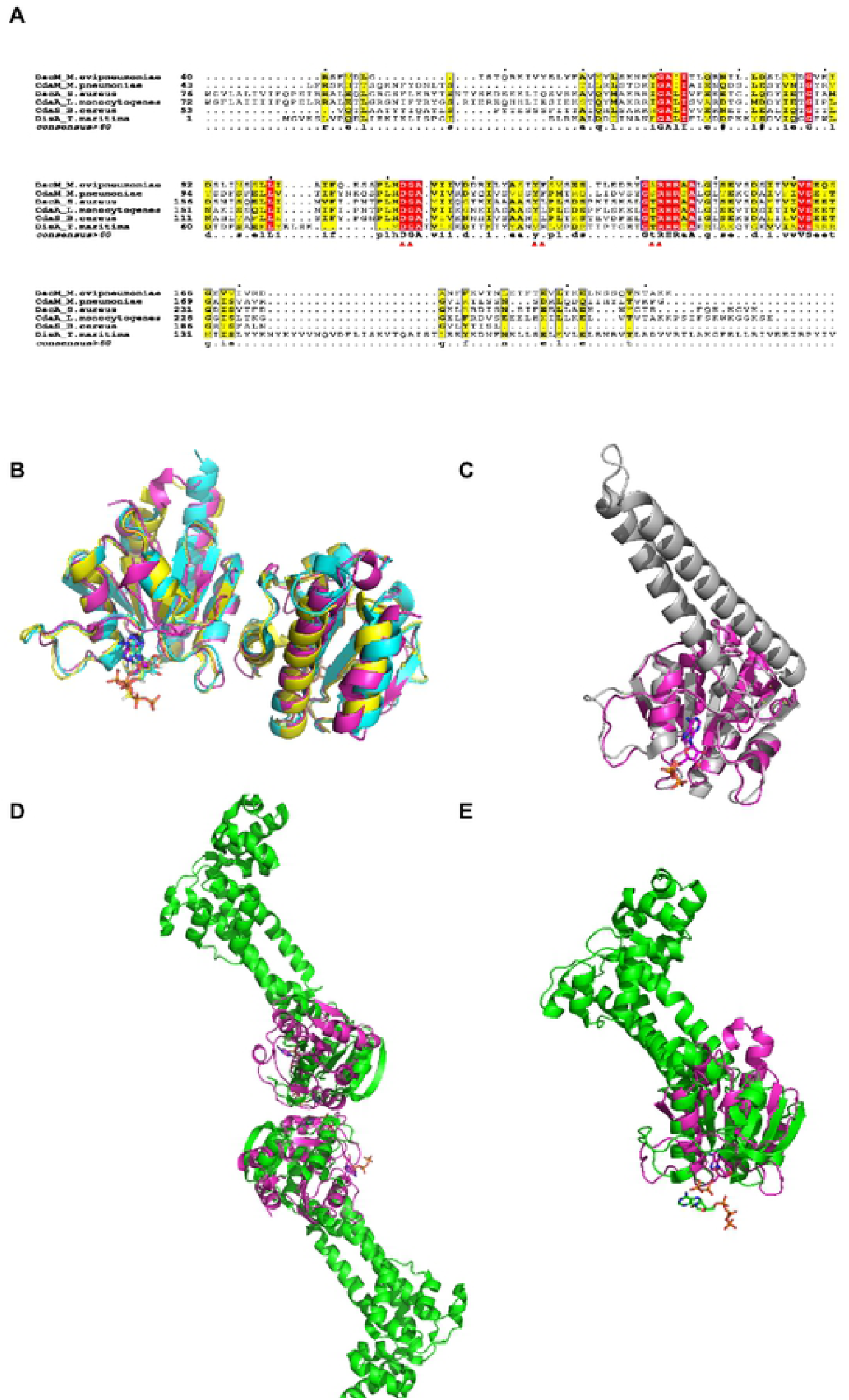
Structure-based alignment of DacM and its homologs from other species. (A) Sequence alignment of DacM (DACs from *M. ovipneumoniae*), CdaM (DACs from *Mycoplasma pneumonia*), DacA (DACs from *Staphylococcus aureus*), CdaA (DACs from *Listeria monocytogenes*), CdaS (DACs from *Bacillus cereus*) and DisA (DACs from *Thermotoga maritima)*. The residues involved in the active site are labeled with red triangles. (B) Structural superimposition of DacM (magenta), DacA (yellow, PDB code 6gyx), and CdaA (blue, PDB code 6hv1) in substrate-bound form. (C) Structural superimposition of one protomer of DacM (magenta) and CdaS (gray, PDB code 2fb5) in substrate-bound form. (D) Structural superimposed of the DacM (magenta) and DisA (green, PDB code 4yvz) dimers in substrate-bound form. (E) Structural superimposition of one protomer of DacM (magenta) and DisA (green, PDB code 4yvz) in substrate-bound form. The substrate molecules, such as ATP, AMP and ApCpp, are shown as sticks.

The structural comparison among these proteins did not reveal major changes, which is consistent with the results of the sequence alignment. As shown in Fig 6B, the DAC domains of DacM, DacA and CdaA are all dimers and present a similar overall fold, overlapping with an rmsd of 0.6 to 0.9 Å. Considering that CdaS is a home-trimer and DisA is a homooctamer, we only compared the protomer of each protein between DacM, CdaS and DisA (Figs 6C and 6E). The DAC domains of DacM and CdaS have an almost identical arrangement with an rmsd of 0.523 Å. By contrast, the DAC domains of DacM and DisA have a slightly different arrangement, with an rmsd of 4.9 Å. However, it should be noted that the orientation of the active-site cavity is almost identical. Additionally, the dimer of DacM and DisA presented a quite different packing orientation (Fig 6D).

## Discussion

In this study, we identified the diadenylate cyclase DacM of *Mycoplasma ovipneumoniae* and confirmed its c-di-AMP synthesis activity in vitro and in *E. coli* cells. Moreover, the apo structure and the structure with two different bound nucleotides were obtained. One remarkable difference among DAC homologs is the number of β-strands. DacM has six mixed β-strands, rather than seven β-strands as observed in its homologs. In fact, the lacking β-strand is replaced by the loop between helices α2 and α3, which also forms the interface of the homodimer. This indicates a different packing mode for DacM, in which more key residues are involved in dimerization (Figs 3C and 6D). Additionally, we obtained the structures of dimer molecules in which the substrate binding cavity was facing outward. However, it is necessary that the two substrate molecules of diadenylate cyclase are placed in face-to-face orientation for c-di-AMP synthesis. This arrangement, called the functional dimer or post-catalytic state, was observed in DisA from *Thermotoga maritima*[23]. Therefore, the substrate-bound structure we solved is thought to be an inactive state, and the two protomers are too far for c-di-AMP synthesis. It is likely that the post-catalytic state of DacM is too transient to capture.

As previously elucidated, few major changes were seen between the apo state and substrate-bound state, except for two notable differences. One is the orientation of Tyr128 in protomer A, which was found to rotate towards the nucleotide when binding ATP via a π-π-stacking interaction. The other is the loop region starting from β3 and near the N-terminal end of α4 helix in protomer B, which makes it more accessible for ADP binding than in the apo state (Figs 4C and 4D). Notably, the latter change makes the DacM dimer molecules more homogeneous. After binding the ADP molecule, protomer B exhibits almost the same arrangement as protomer A in the apo state and substrate-bound state (S5C Fig). This is likely a reason why the substrate-bound crystal exhibited better diffraction. The active-site cavity of DAC proteins is highly conserved. In particular, the Tyr exhibits an unequivocally conserved role in the ATP-binding process, and mutation of this site significantly reduced DacM catalytic activity. Mutations at other active sites, such as Asp112, Gly113, Phe129, and Arg143 were lethal to DacM catalytic activity, its catalytic activity is significantly reduced or completely lost (Figs 5A and 5B). Among these residues, Phe129 draw our attention because its employ of enzymatic activity has not been reported in its homologues yet. Other residues, Asp112, Gly113, Tyr128 and Arg143 are conserved with homologues in enzymatic activity as reported (17). By analysis combined with the structure-based sequence alignment between DacM and its homologues from other species, which Phe129 is not strict conservation (Fig. 6A), it may reveal a unique or more complicated substrate binding mechanism of DacM.

It should be noted that the enzyme activity of the other DAC proteins requires mental ions, such as Mg^2+^, Mn^2+^ or Co^2+^. However, our present work is limited to discuss the metal ion dependence in details. This question still awaits more structural and experimental investigation in future. Taken together, our work provides a basis for the rational design for novel antibiotics that can be used in animal husbandry. Nevertheless, additional efforts are needed to elucidate the entire picture of c-di-AMP signaling and better understand the underlying mechanisms.

## Materials and methods

Materials and detailed protocols are described in *Supporting Information*

### Isolation and detection of c-di-AMP from *M*.*ovipneumoniae*

The c-di-AMP was isolated from *M. ovipneumoniae* Y98 and detected by LC-MS[26, 27] on an ACQUITY UPLC I-Class system (Waters, UK) with a VION IMS QTOF Mass spectrometer (Waters, UK). A commercial standard of c-di-AMP (InvivoGen, Hong Kong, China) was used to determine the concentration by comparing the peak areas.

### Plasmid Construction

E. coli BL21 (DE3) and plasmid pET-28b (+) were used for the expression of recombinant diadenylate cyclase protein. The dacM sequence was codon-optimized for E. coli and synthesized by GenScript Corporation (Nanjing, China).

### Analysis of enzyme activity in vivo and vitro

To assess the biochemical activity of candidate diadenylate cyclases in vivo, the enzymes were expressed in *E. coli* BL21(DE3), which does not produce c-di-AMP[25]. The c-di-AMP was detected by LC-MS.

The activity of diadenylate cyclase in vitro was measured by detecting the c-di-AMP production. Briefly, the reaction was started by adding 2 μM Δ^2-33^DacM to reaction mixture containing 50 mM Tris-HCl pH 7.5, 10 mM MgCl_2_, 150 mM NaCl, and 0.1 mM ATP or 0.1 mM ADP. The c-di-AMP was detected by LC-MS[27] after reaction mixture incubated for 1 h at 37 °C. All activity measurements were carried out in triplicate. Details are given in SI Materials and Methods.

### Protein purification and crystallization

A truncated version of the *dacM* gene (Δ^2-33^DacM) was subcloned into the pET28b vector. The recombinant protein was overexpressed in *E. coli* BL21 (DE3) and purified by nickel affinity column, followed by purification to homogeneity by gel-filtration chromatography (Superdex 75 16/600, GE Healthcare) Both apo and substrate-bound proteins were crystallized at 16 °C using the hanging-drop vapor-diffusion method. Crystals of apo-DacM and the complex with good diffraction were grown in the reservoir solution containing 0.1 M citric acid pH 3.5, 1 M sodium malonate pH 5.0 and 0.1 M HEPES pH 7.5 with 19% (w/v) PEG 3350, respectively.

### X-ray data collection and structure determination

All native datasets were collected at the Shanghai Synchrotron Radiation Facility beamline BL17U and were processed using HKL2000[28] in conjunction with programs from the CCP4 suite[29]. The initial phases were obtained using the molecular replacement method in PHASER[30] with the structure of CdaA from *Listeria monocytogenes* (PDB ID: 4RV7) as the search model. The models of apo form and substrate-bound form were built and manually refined using COOT[31] and PHENIX[32], respectively.

### Analysis of DacM mutants activity

In this experiment, we mutated the Asp112, Gly113, Tyr128, Phe129, and Arg143 sites of DacM to Ala for gene synthesis by Corebiolab (Wuhan, China). The mutant genes were cloned into pET-28b (+) expression vector, and transformed into E. coli BL21(DE3) strain, picked and sequenced. The successfully DacM mutants was cultured at 37°C for 12 hours, c-di-AMP was extracted, and the DacM mutants catalytic activity was examined in vivo by LC-MS.

## Acknowledgments

This work was supported by the National Natural Science Foundation of China (31960036), Key R & D projects of Ningxia Hui Autonomous Region (2017BN04)

## Supporting information

### Supporting information text (Materials and detailed protocols)

**S1 Fig. Bioinformatics analysis of DacM**. (A) Partial sequence alignment of DacM from *M. ovipneumoniae* with three other diadenylate cyclases, showing the conserved DGA and RHR motifs. (B) Schematic representation of the primary structure of DacM. TM, transmembrane region; DAC, diadenylate cyclase domain; #, DGA active site motif; *, RHR active site motif.

**S2 Fig. Purification and western blot analysis of recombinant DacM**. (A) Fractions of the purification of recombinant DacM on a Coomassie blue-stained SDS-PAGE (12%). MW: molecular weight ladder, Lane 1: Purified recombinant DacM. (B) Western blot analysis using polyclonal antibodies against DacM. MW: molecular weight ladder, Lane 1: western blot detection of recombinant DacM.

**S3 Fig. Immunogold labeling of DacM protein in cells of *M. ovipneumoniae* Y98 strain**. (A) Cells of *M. ovipneumoniae* without antibodies as negative control. (B) *M. ovipneumoniae* incubated with anti-DacM polyclonal antibodies.

**S4 Fig. Western blot analysis of the membrane protein fraction of *M. ovipneumoniae* strain Y98 DacM using polyclonal antibodies against DacM**.

**S5 Fig. Detailed structural comparison of Δ**^**2-33**^**DacM**. (A) Comparison between protomer A (colored pink) and protomer B (colored light green) of the DacM apo form. (B) Comparison between protomer A (colored magenta) and protomer B (colored cyan) of the DacM substrate-bound form. (C) Comparison between protomer A of the DacM apo form and protomer B of the DacM substrate-bound form. The key residue Tyr128, whose side chain is shown as sticks in each protomer, is colored orange (protomer A of the DacM apo form), green (protomer B of the DacM apo form), red (protomer A of the DacM substrate-bound form) and dark blue (protomer B of the DacM substrate-bound form). The ATP molecule is colored yellow and the ADP molecule is colored gray.

**S1 Table. Data collection and refinement statistics**.

## Author Contributions

**Conceptualization:** Xiujing Hao, Xinhui Zhou, Shilong Fan, Min Li.

**Formal analysis:** Xiujing Hao, Xinhui Zhou, Shilong Fan, Min Li.

**Data curation:** Xiujing Hao, Xinhui Zhou, Ying Zhang, Yang Han, Zhaokun Xu, Chunji Ma,

Haixia Luo, Kemin Qi

**Funding acquisition:** Xiujing Hao, Xinhui Zhou, Shilong Fan, Min Li.

**Investigation:** Ying Zhang, Yang Han, Zhaokun Xu, Chunji Ma, Haixia Luo, Kemin Qi.

**Methodology:** Xiujing Hao, Xinhui Zhou, Chunji Ma, Haixia Luo, Kemin Qi

**Project administration:** Shilong Fan, Min Li.

**Supervision:** Shilong Fan, Min Li.

**Writing – original draft:** Xiujing Hao, Xinhui Zhou, Ying Zhang.

**Writing – review & editing:** Xiujing Hao, Xinhui Zhou, Ying Zhang.

## Competing Interest Statement

The authors declare no conflict of interest.

